# Reversible block of cerebellar outflow reveals cortical circuitry for motor coordination

**DOI:** 10.1101/429043

**Authors:** Abdulraheem Nashef, Oren Cohen, Ran Harel, Zvi Israel, Yifat Prut

**Affiliations:** Dept. of Medical Neurobiology, IMRIC and ELSC, The Hebrew University, Hadassah Medical School, Jerusalem, 9112102, Israel; Dept. of Neurosurgery, Sheba Medical Center, Tel Aviv, Israel; Dept. of Neurosurgery, Hadassah Hospital, Jerusalem, Israel

**Keywords:** cerebellar-thalamo-cortical, high-frequency stimulation, cerebellar ataxia, non-human primates, noise correlation, motor timing, inter-joint coordination

## Abstract

Coordinated movements are achieved by selecting muscles and activating them at specific times. This process relies on intact cerebellar circuitry, as demonstrated by motor impairments triggered by cerebellar lesions. Based on anatomical connectivity and symptoms observed in cerebellar patients, we hypothesized that cerebellar dysfunction should disrupt the temporal patterns of motor cortical activity but not the selected motor plan. To test this hypothesis, we reversibly blocked cerebellar outflow in primates while monitoring motor behavior and neural activity. This manipulation replicated the impaired motor timing and coordination characteristic of cerebellar ataxia. We found extensive changes in motor cortical activity, including a loss of response transients at movement onset and a decoupling of task-related activity. Nonetheless, the spatial tuning of cells was unaffected and their early preparatory activity was mostly intact. These results indicate that the timing of actions, but not the selection of muscles, is regulated through cerebellar control of motor cortical activity.

**HIGHLIGHTS:**

- High frequency stimulation blocked cerebellar outflow and impaired motor behavior
- Response patterns and coordinated firing of CTC neurons were disrupted
- The spatial tuning and early preparatory activity of neurons were unaffected
- Cerebellar control of local and global cortical synchrony supports motor timing

**IN BRIEF:** Nashef et al. used high frequency stimulation to block cerebellar outflow. This manipulation impaired motor timing and coordination similarly to symptoms found in cerebellar patients. In parallel, the response patterns of cortical neurons and cell-to-cell synchronization were altered, yet spatial tuning was maintained. Motor timing and coordination are regulated by a dedicated cerebellar signal that organizes execution-related activity of a motor cortical subnetwork.

## INTRODUCTION

In daily life, well-coordinated and properly timed movements are performed in an effortless manner. This ability is considered to be in large part mediated by cerebellar shaping of motor output (Bastian et al., 1996; Beaubaton et al., 1978; Holmes, 1939; Machado et al., 2015; Meyer-Lohmann et al., 1977; Schlerf et al., 2007; Spencer et al., 2003). This claim is based on studies of motor behavior in cerebellar patients (Bastian et al., 1996; Bo et al., 2008; Spencer et al., 2003) which indicate that these subjects suffer from poor timing of actions (Schlerf et al., 2007; Spencer et al., 2003) and tend to produce abnormally curved and uncoordinated movements (Bastian et al., 1996). However, the neural mechanisms through which the cerebellum controls these motor functions are controlled by the cerebellum are still unclear.

Cerebellar impact on voluntary movements of the upper limb is predominantly mediated by two pathways. The cerebellar-rubro-spinal tract provides the cerebellum with fast access to segmental circuitry (Cohen et al., 2017; Garwicz, 2002; Huisman et al., 1983; Nioche et al., 2009) but its importance diminished considerably over the course of evolution (Nathan and Smith, 1982; Padel et al., 1981; Schoen, 1964; ten Donkelaar 1988). By contrast, the cerebellar-thalamo-cortical (CTC) pathway (Rispal-Padel et al., 1981) increased in size with evolution and in primates became the dominant route in mediating the cerebellar control of voluntary movements (Horne and Butler, 1995). The CTC pathway originates in the deep cerebellar nuclei, primarily in the dentate nucleus (Wiesendanger and Wiesendanger, 1985), makes remarkably effective synaptic contacts with the cerebellar-receiving areas of the motor thalamus (Asanuma et al., 1983a; Asanuma et al., 1983b; Aumann et al., 1994; Sakai et al., 1996; Shinoda et al., 1982) and terminates extensively throughout the motor cortex, creating patches of terminations which may extend several millimeters in the rostrocaudal axis (Shinoda et al., 1993). Cooling the deep cerebellar nuclei in primate models was shown to trigger behavioral symptoms similar to those found in cerebellar patients and concomitant changes in motor cortical activity (Hore and Flament, 1988; Meyer-Lohmann et al., 1975), most of which involved decreased activity at movement onset. These results indicate that the CTC system has online access to evolving motor commands and thereby affects motor actions. However, it remains unclear what features and parameters of the motor command are specifically dictated by the CTC system, and in what way the loss of the CTC drive, which apparently constitutes only a small fraction of the input to motor cortical neurons (Bopp et al., 2017), affects the firing of single cells thus leading to impaired timing and coordination across multiple effectors.

To address these questions, we trained two monkeys to perform a center-out reaching task which relied on predictive timing (Bares et al., 2007; Bo et al., 2008). Stimulating electrodes were chronically implanted in the superior cerebellar peduncle (SCP) and recordings were made simultaneously from multiple cortical sites. Single-pulse stimulation was used to identify motor cortical neurons that are part of the CTC pathway, and high-frequency stimulation (Agnesi et al., 2013; Agnesi et al., 2015; Chiken and Nambu, 2016; Dostrovsky and Lozano, 2002) was used to interfere with the normal flow of information through the pathway. Using this method, we identified a large number of motor cortical neurons that were part of the CTC system. High frequency stimulation effectively prevented information flow in the CTC pathway and produced reversible motor deficits similar to those found in cerebellar patients, including impaired timing and coordination of movements. The observed behavioral deficits were preceded by substantial changes in neural activity. Specifically, cortical cells that were part of the CTC system expressed delayed and sluggish response onset, but their spatial tuning was unaffected. Changes in the response pattern were confined to time of movement execution without affecting early stages of motor preparation. In addition, the signal-dependent noise correlation typically found between neighboring motor cortical cells was lost. Finally, the well-organized recruitment order of cells that were recorded from shoulder and elbow-related cortical sites was disturbed. All these changes in neural activity and behavioral deficits reversed back to baseline as soon as the stimulation was stopped. These results suggest that the cerebellar impact on motor cortical activity is not limited to regulating single cell activity; rather CTC input acts to locally synchronize and temporally organize the activity of a spatially distributed motor subnetwork. However, CTC regulation of temporal properties of motor cortical firing operates independently of the mechanism dictating spatial tuning. The local and global synchrony triggered by the CTC system can enhance the throughput of cortical units and control motor timing and coordination.

## RESULTS

We recorded neural activity from the sensorimotor cortex (**Fig. 1A**) in response to single-pulse SCP stimulation and while the monkeys performed a center-out reaching task (**Fig. 1B**). Figure 1C presents the recording maps obtained for the two monkeys (right and left hemispheres in monkey C and right hemisphere in monkey M). A high proportion of motor cortical sites across the entire recording area showed a significant multiunit response to stimulation (68%-73% between monkeys). During task performance, we recorded neural activity from multiple single cells and measured their task-related activity (**Fig. 1D)** and response to SCP stimulation (**Fig. 1E**). The onset time and response pattern expressed by cortical cells in response to SCP stimulation was similar to our previous findings (Nashef et al., 2018) and consistent with the di-synaptic impact of SCP stimulation on cortical cells.

**Figure 1.**
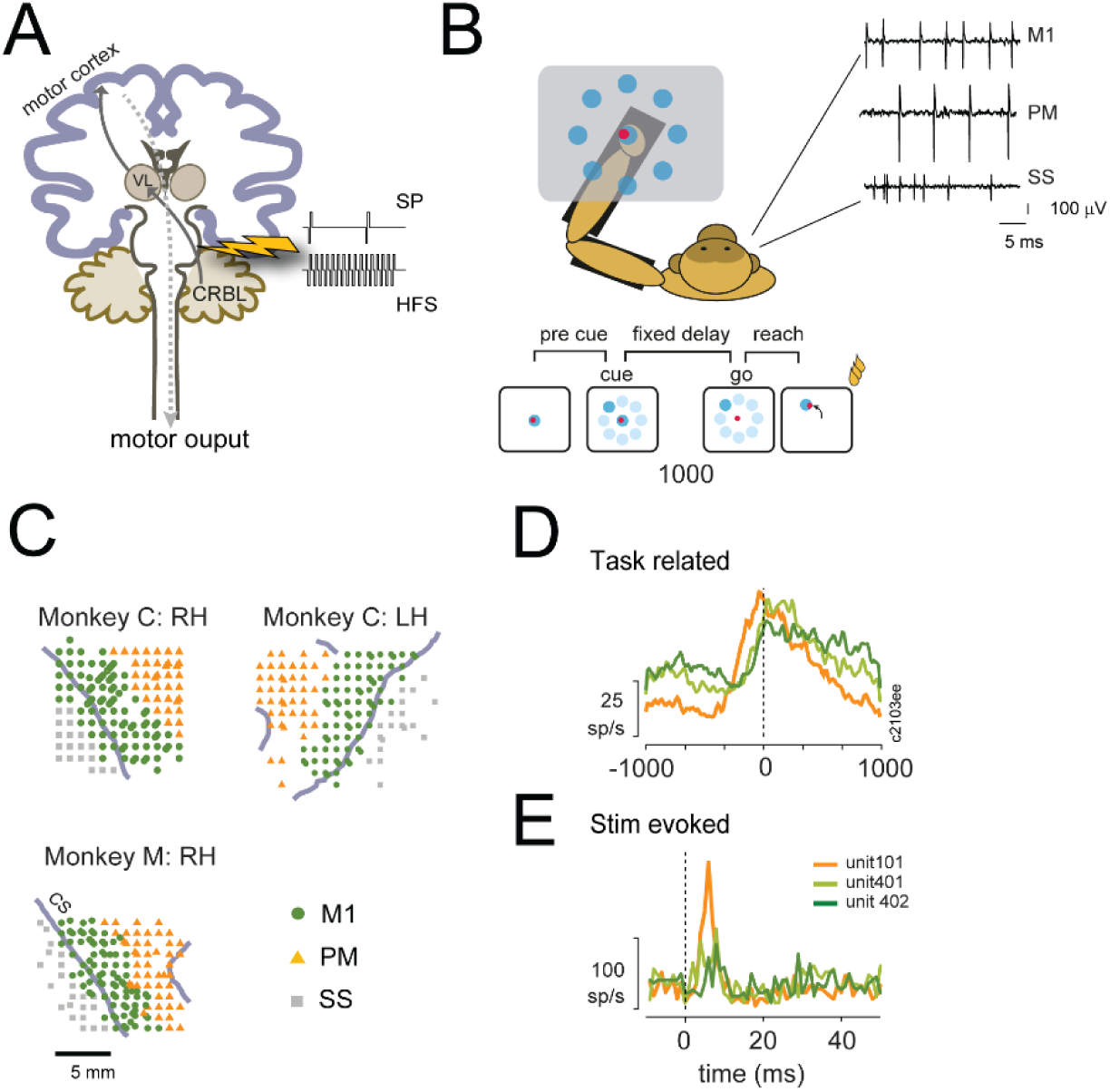
Experimental design and recording configuration. (**A**) A schematic view of the CTC system. The system originates from cells in the deep cerebellar nuclei (primarily the dentate nucleus). Axons of these cells make initial synaptic contact in the motor thalamus, mainly the ventrolateral nuclei (VL). Thalamocortical fibers make a second synaptic contact in the motor cortex. Motor cortical output projects downstream to primarily affect the contralateral part of the body (ipsilateral to the cerebellar projection site). Neural activity was recorded from the motor and somatosensory areas of the cortex while stimulations were applied to the superior cerebellar peduncle (SCP) according to one of two stimulation protocols: single pulse (SP) bipolar stimulation applied at low frequency or high frequency stimulation (HFS). CRBL-cerebellum. (**B**) Behavioral paradigm. Monkeys were trained to wear an exoskeleton and control a cursor that appeared on a horizontally positioned screen. The movement of the monkeys was constrained to planar, shoulder-elbow reaching movements. The sequence of events composing a single trial included a pre-cue period, target onset (where 1 of 8 equally distributed targets appeared), a delay period and a “go” signal, after which the monkey had to acquire the target within a pre-defined movement time. Correct performance resulted in a reward (a drop of applesauce). Neural activity was recorded simultaneously from the motor and sensorimotor cortical areas. M1-primary motor; PM-premotor; SS-somatosensory. Vertical scale bar: 100 μV; Horizontal scale bar: 5 ms (**C**) Cortical maps obtained from monkey C (right and left hemispheres: RH, LH) and monkey M (right hemisphere only). CS-central sulcus. Scale bar- 5 mm (**D**) Peri-event time histogram calculated for three, simultaneously recorded cortical neurons around movement onset (t=0). Two neurons were recorded from M1 (green curves) and one from the PM cortex. (**E**) Evoked response of the three neurons shown in (D) to single pulse SCP stimulation.

### High frequency SCP stimulation interrupts the transmission of information in the CTC pathway

Studying the cerebellar impact on motor cortical activity can benefit from comparing neural activity obtained when blocking the flow of information in the CTC pathway to the baseline level. Previous studies have used dentate cooling to identify cerebellar involvement in motor cortical activity and motor behavior (Flament and Hore, 1986; Meyer-Lohmann et al., 1975). However, this approach cannot dissect the underlying circuity that mediates the observed deficits. In addition, it is difficult to estimate the efficiency of this manipulation in blocking information flow through the CTC pathway. Instead, we implemented the commonly used high frequency stimulation protocol that was shown to interfere with ongoing, often pathological patterns of neural activity (Benabid et al., 1991; Hodaie et al., 2002; Limousin et al., 1998; Torres et al., 2010). We applied the same stimulation protocol through the SCP electrode and relied on the fact that synaptic transmission cannot follow this high frequency activation pattern (Agnesi et al., 2015; Iremonger et al., 2006; Wang and Kaczmarek, 1998; Zucker and Regehr, 2002). To verify the negative effect of HFS on information transfer, we tested the changes in neural response to SCP stimulation when using HFS as compared to low frequency stimulation. Figure 2A presents an example of a neural response to single-pulse SCP stimulation exhibited by a cortical cell. In this example, the stimulation triggered a response in a large fraction of the sweeps. However, when the same stimulus was applied at a higher frequency (**Fig. 2B)**, the tight contingency between the stimulus and the single cell response was lost. This change in response to stimulation is clearly apparent in the raster plot and the peri-stimulus time histogram (PSTH) computed for that cell (**Fig. 2C**). Across the population, SCP-responsive cells had a significantly weaker response to stimuli applied during high frequency stimulation than to a single pulse stimulation applied at the same intensity level. This is shown in the stimulus-triggered mean rate for responsive cortical cells (**Fig. 2D**) for single pulse (black trace), HFS (red trace) and the baseline level computed using artificially injected “stimulation times” in control trials (surrogate - blue trace). Computing the baseline level in this manner ensured that both rate measures (baseline and post-stimulus) were similarly affected by the task-related rate modulation. We further quantified this reduction in response magnitude by calculating the firing rate during the 6 ms following the stimulation pulses (for single pulse stimulation, HFS protocol, and control trials). As expected, during single-pulse stimulation, the firing rate of the cortical cells increased compared to the baseline level (**Fig. 2E**). This was expected since we only tested cells that were responsive to SCP stimulation. However, during HFS, the post-stimulus firing rate of responsive cells declined considerably compared to the single pulse stimulation (Wilcoxon signed-rank; p<0.001) even though we used the same stimulus intensity and only changed the stimulation frequency. In fact, the post-stimulus response area (i.e., the number of additional spikes triggered by the stimulus) for cortical cells dropped to 86.4% during HFS compared to the single pulse stimulation. Further, during the HFS trials, the post-stimulus rate of the cortical cells was similar to the baseline rate computed when no stimulation was applied (**Fig. 2E**, p<0.49). Finally, in a separate study conducted on a third monkey we tested the frequency-dependency of response suppression and found that it occurred at frequencies exceeding 30 Hz (**Fig. S1**). This result is consistent with findings obtained *in vitro* (Gornati et al., 2018) showing that the transmission between the cerebellar and motor thalamus is suppressed during high frequency stimulation. We argue that the observed response suppression during HFS can be considered as a practical and efficient block of the CTC pathway. This conclusion is based on the argument that a single SCP stimulation pulse can be viewed as a highly-synchronous packet of spikes that propagates through the SCP to the motor thalamus. This synchronous activation contrasts with the innate activation of the system which is expected to be relatively sparse, slow and less efficient in triggering a post-synaptic response. Therefore, the pronounced drop in single cell response to the highly potent SCP activation provides an upper bound on the actual information transfer through the pathway during this time. Taken together, these results suggest that HFS efficiently blocked the flow of information from the cerebellar output to the motor cortex.

**Figure 2.**
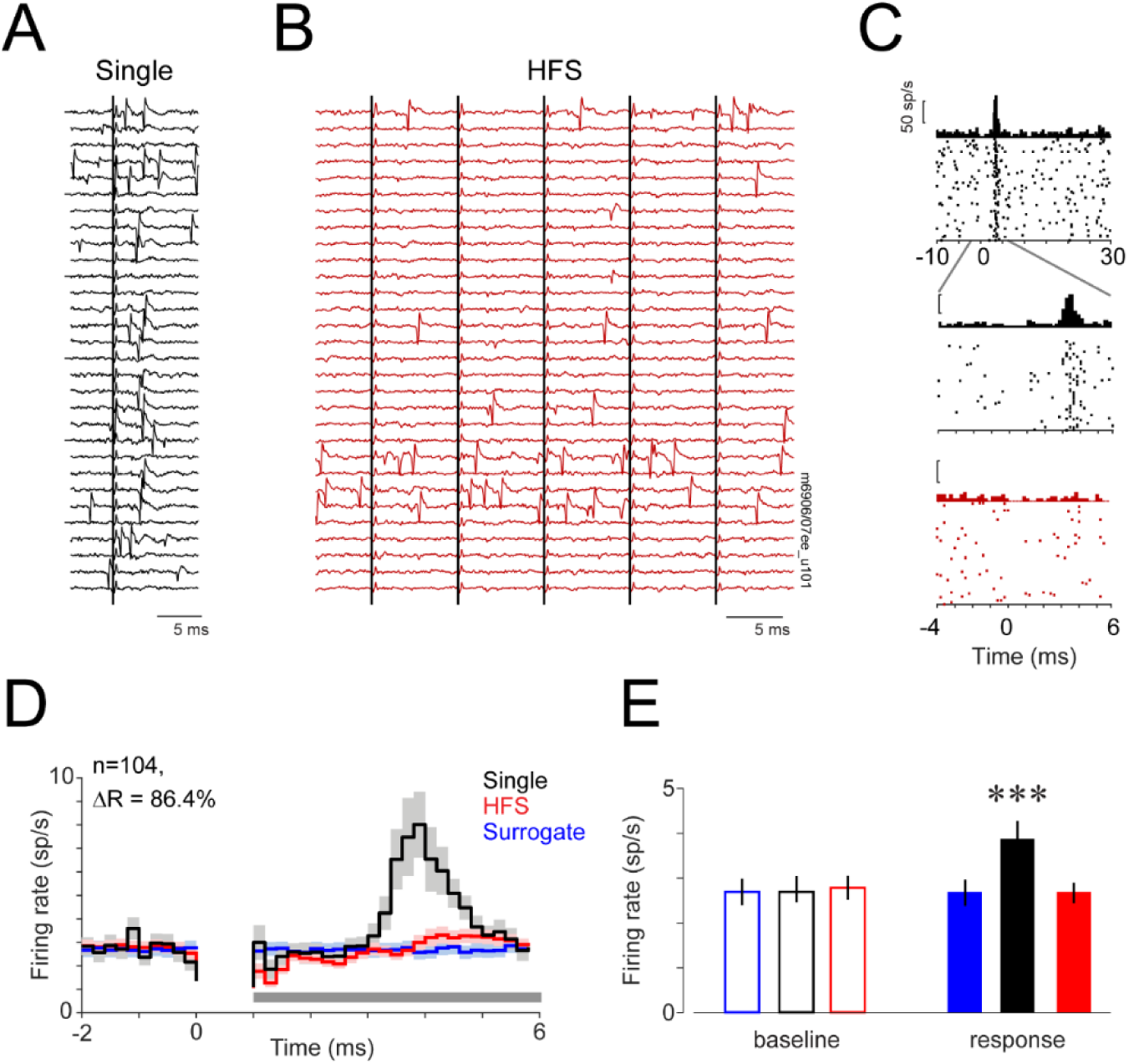
Neural response to SCP stimulation was abolished at high frequencies. (**A**) Example of a single neuron response to SCP stimulation applied at low frequency (single-pulse stimuli). The figure shows 30 randomly selected stimulation sweeps. The single sweep response of the cell is variable but clearly observable. Scale bar: 5 ms (**B**) Activity of the same cortical cell during stimulation applied at the same intensity as in (A) but at high frequency (HFS). Each trace captures 5 successive stimuli. In this case, successive traces were obtained at regular intervals along the entire HFS epoch. Upper traces present early occurring stimuli during the HFS, whereas bottom traces are from late parts of the HFS. Scale bar: 5 ms. (**C**) Raster plots and PSTHs of the single cell response to SCP stimulation. Same neuron as shown in (A) and (B). Top plot, response of cells to single pulse stimulation. Middle plot shows the same single-pulse response but at an extended time scale. Bottom plot presents the response of the same neuron to stimulation applied at high frequency. Scale bars: 50 sp/s. (**D**) Average cortical response computed by averaging the single cell response (quantified by the average firing rate around stimulus onset time) across all responsive cortical neurons (n=104) for single pulse stimulation (single, black), high-frequency stimulation (HFS, red) and control (surrogate, blue). Single cell responses were computed using a 0.2 ms bin size. The gap after time zero reflects the stimulus-related dead-time in spike detection. Shading around each response curve depicts the standard error of the mean (**E**) Comparisons of post-stimulus firing rate, computed between 1 and 6 ms after stimulus onset to pre-stimulation baseline (defined between −2 to 0 ms) for the three protocols (surrogate control, single pulse and HFS). For each epoch we compared the post-stimulation rate (full bars) to the pre-stimulation baseline level (empty bars) using Wilcoxon’s signed-rank (***, p < 0.001). **See also Fig. S1**.

### High-frequency stimulation alters timing and coordination of reaching movements

We found that motor behavior was modified considerably during HFS trials. Figure 3A shows an example of a single recording session in which hand trajectories became more variable during HFS trials than in control trials. We quantified the change in motor performance using the response and movement times computed for each trial based on the trajectory (**Fig. 3B**). For this analysis we only considered correctly performed trials and since task design encouraged the monkey to predict the onset time of the go signal, the response time (RT - time between “Go” signal and movement onset) was often negative. During HFS, the mean trajectory across all sessions was longer and more variable, as seen in the plots for the mean and standard deviation of the center-to-target movement traces (**Fig. 3C**). Response time increased significantly (from −183.3 ms during control to - 133.1 ms during HFS, paired t-test, p<0.001), as did movement time (from 446.9 ms to 517.3 ms, paired t-test, p<0.001); the path length was longer (from 4.19 cm to 4.53 cm; p<0.02), and velocity decreased (from 10.34 cm/s to 9.74 cm/s; p<0.001) for trials performed during HFS (**Fig. 3D**). These changes in motor behavior reversed back to the baseline level when HFS was halted (washout bars in **Fig. 3D**). In contrast to the effect of HFS on motor behavior, the single pulse stimulation protocol had no effect on motor behavior despite its impact on many motor cortical cells (**Fig. 3D**, black bars). These findings suggest that when stimuli are applied at low frequencies, the brief overt activation caused by SCP stimulation is insufficient to alter motor behavior; rather, the temporary block of information flow in the CTC pathway that takes place during high frequency stimulation is responsible for the observed effects.

**Figure 3.**
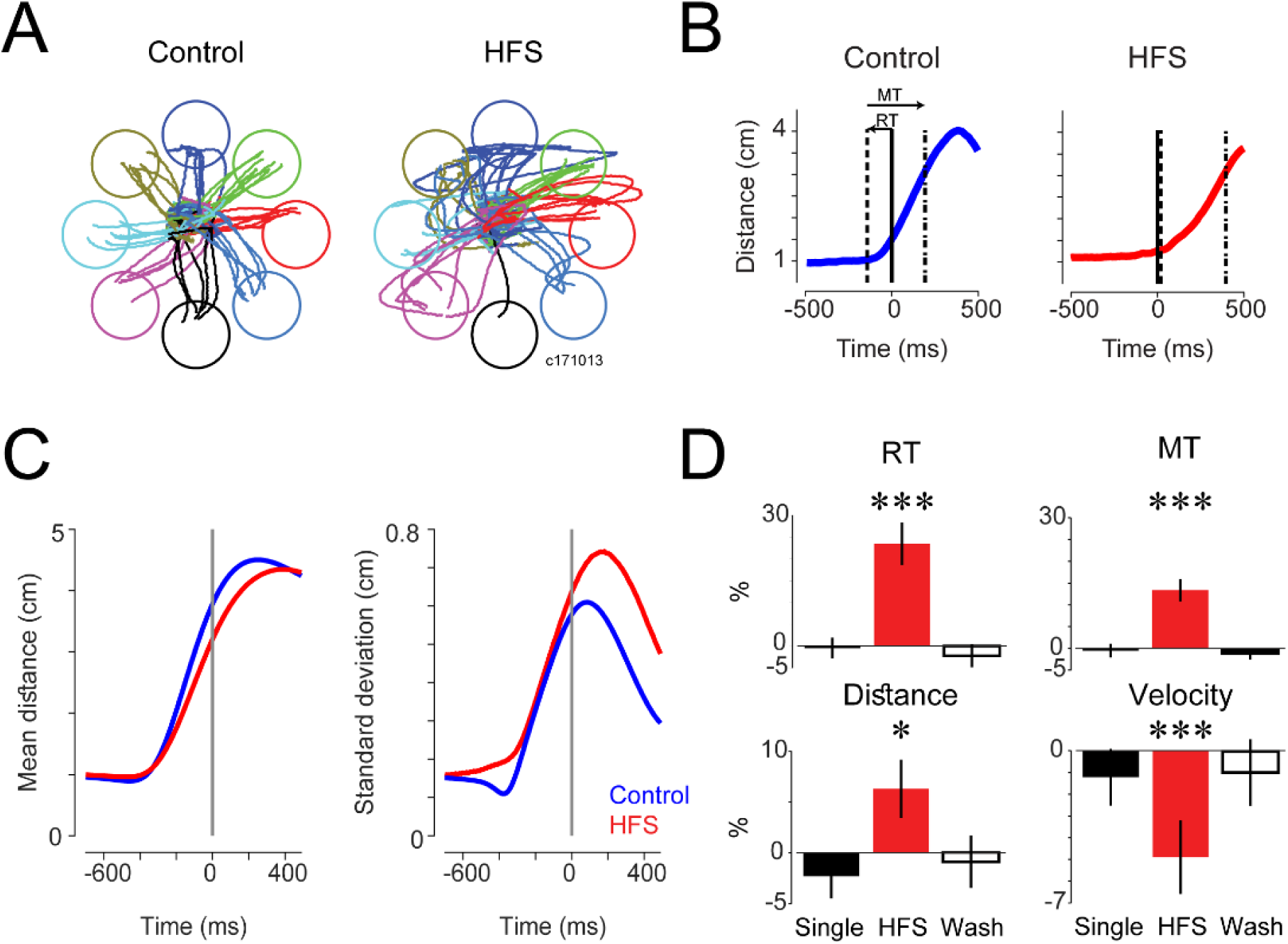
Motor behavior was impaired during high frequency stimulation. (**A**) Example of a single recording session showing the effect of HFS on movement trajectory during control (left) and HFS trials. Each plot depicts 50 trials. (**B**) Single trial example of the radial distance between the cursor and the center target computed during two trial conditions: control (left) and HFS (right). In this example, both movements were made to the same peripheral target. The radial distance trace was used to calculate the single trial response time (RT; for control: −92.2 ms and for HFS: 18.9 ms) and the movement time (MT; control: 284.3 ms, HFS: 375.4 ms; see Methods). (**C**) Average (left) and standard deviation (right) of the radial distance computed for control (blue) and HFS (red) trials and averaged over 110 recording sessions. (**D**) Modulation of behavioral parameters computed during single pulse stimulation (black bars), HFS (red bars) and post-HFS (washout) trials (empty bars). Parameters include RT, MT, path distance and average movement velocity. Changes in parameters are shown in percentages relative to the baseline level (computed during control trials). In all cases, the parameter was significantly modulated solely during HFS. Significance levels were evaluated using paired t-tests. *** indicates significance at p < 0.001; * indicates p < 0.05.

Previous studies have suggested that cerebellar involvement is particularly important in controlling inter-joint coordination (Bastian et al., 1996; Goodkin et al., 1993; Holmes, 1939; Thach et al., 1993; Thach et al., 1992). We therefore tested the effect of HFS on the elbow-to-shoulder coordination required to perform the behavioral task used in this study. We took advantage of the exoskeleton system (Scott, 1999) worn by the monkeys that continuously recorded their elbow and shoulder motions. Previous studies have reported that cerebellar patients tend to exhibit curved hand trajectories (Deuschl et al., 2000; Martin et al., 2000). We computed the curvature index (Deuschl et al., 2000), which quantifies the average distance of the trajectory from a straight line connecting the same start and end points. During HFS trials the curvature index increased significantly compared to the control trials (**Fig. 4A**; p<10^−10^, Wilcoxon’s signed-rank). These results show that applying HFS further replicates deficits in motor coordination similar to those shown in cerebellar patients, in a reversible manner.

**Figure 4.**
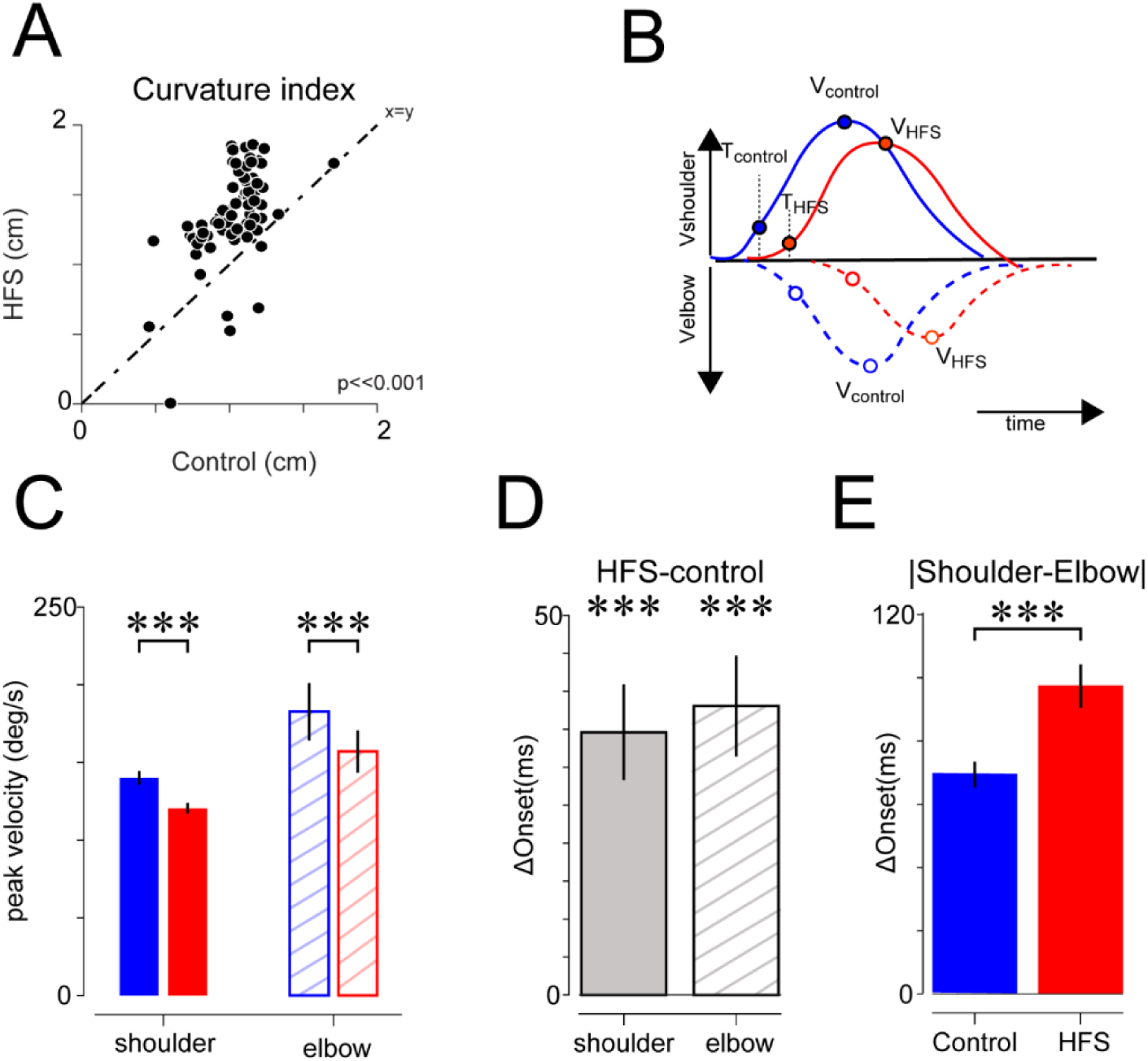
HFS impaired inter-joint coordination. (**A**) Relations between the curvature index (see Methods) computed during control and HFS calculated during movement time (paired t-test, p<10^−8^; n=77 recording sessions). (**B**) Schematic illustration of a shoulder (solid line) and elbow (dashed) angular velocity profile recorded during control (blue) and HFS (red) trials for a movement to a specific target. Using the velocity profile we calculated the maximal velocity (V_control_, V_HFS_) and movement onset time (T_control_, T_HFS_) defined as the time when velocity exceeded a threshold value of 0.5 × baseline noise for a consistent time period (see Methods). (**C**) Peak angular velocity for shoulder (solid bars) and elbow (hatched bars) joints during control (blue) and HFS (red) trials (paired t-test, p<0.001 for both joints). (**D**) Mean change in onset time for shoulder (solid) and elbow (hatched) joints during HFS application relative to onset time computed during control trials (paired t-test, p<0.001). (**E**) Mean changes in the shoulder to elbow onset latency following HFS application (paired t-test; p<0.001). All values were computed relative to the “GO” signal. **See also Fig. S2**.

Next, we measured changes in movement kinematics between the control and HFS trials. Figure 4B illustrates shoulder and elbow velocity profiles recorded in a single control (solid lines) and HFS (dashed line) trials that were directed to a single target and the kinematic parameters used to quantify these movements. We found that during the HFS trials, both the shoulder and elbow peak velocity decreased significantly (**Fig. 4C**; Shoulder: 140.1±4.47 °/s to 120.5±3.33 °/s, paired t-test: p<0.001; Elbow: 182.7±18.51 °/s to 157±13.6 °/s, paired t-test: p<0.001). Further, the onset time of the shoulder and elbow movements increased significantly during HFS trials (**Fig. 4D** and **Fig. S2** shoulder: 34.6±6.3 ms increase, paired t-test, p<0.001; elbow: 38.1±6.6 ms increase, p<0.001). Finally, the difference between the onset times of the two joints increased as well (**Fig. 4E** and **Fig. S2;** 69.4±4.1 ms to 97.4±6.9 ms, p<0.001), suggesting that during HFS trials, not only was movement onset delayed but also the shoulder and elbow joints tended to be activated in a more isolated manner, as often found in ataxic patients (Bastian et al., 1996; Becker et al., 1990). These results suggest that motor behavior in HFS trials exhibits similar impairments as found in cerebellar patients, further supporting the efficiency of the HFS protocol in blocking cerebellar outflow.

### HFS modifies the movement-related activity of CTC neurons in a manner correlated with and predictive of behavioral changes

After verifying that HFS prevented the normal flow of CTC information and induced considerable changes in motor behavior, we examined the changes in neuronal activity which occurred at the same time and inspected their role in mediating the behavioral impairments. This was done by comparing the task-related activity of cortical neurons that were part of the CTC system during HFS and the control trials. CTC neurons were defined based on their significant, excitatory responses to SCP stimulation (Nashef et al., 2018). Figure 5 presents the activity of one such cell (**Fig 5A**) and its preferred direction (**Fig. 5B**) during control (blue) and HFS (red) trials. The cell was identified as part of the CTC system based on its early excitatory response to SCP stimulation (see inset in **Fig 5A**). In this example, the movement-related activity of the neuron during HFS trials lacked the transient firing at movement onset that occurred during control trials. Despite the change in the response profile, the preferred direction of the cell remained the same. Across the population, cells that were both responsive to SCP stimulation and directionally tuned during the control trials (n=57) expressed a consistent tendency to exhibit more sluggish response profiles during HFS trials (**Fig. 5C**). The weaker activation was not an outcome of a general decrease in the neuronal firing rate during HFS trials since the pre-cue firing of the cells was not significantly different between the HFS and control trials **(paired t-test, p=0.24**). In addition, despite the change in response profile of neurons, there was no significant change in the PD between HFS and the control trials (**Fig. 5D**, mean ΔPD = 0.24 rad, p<0.59, one-sample test for mean direction).

**Figure 5.**
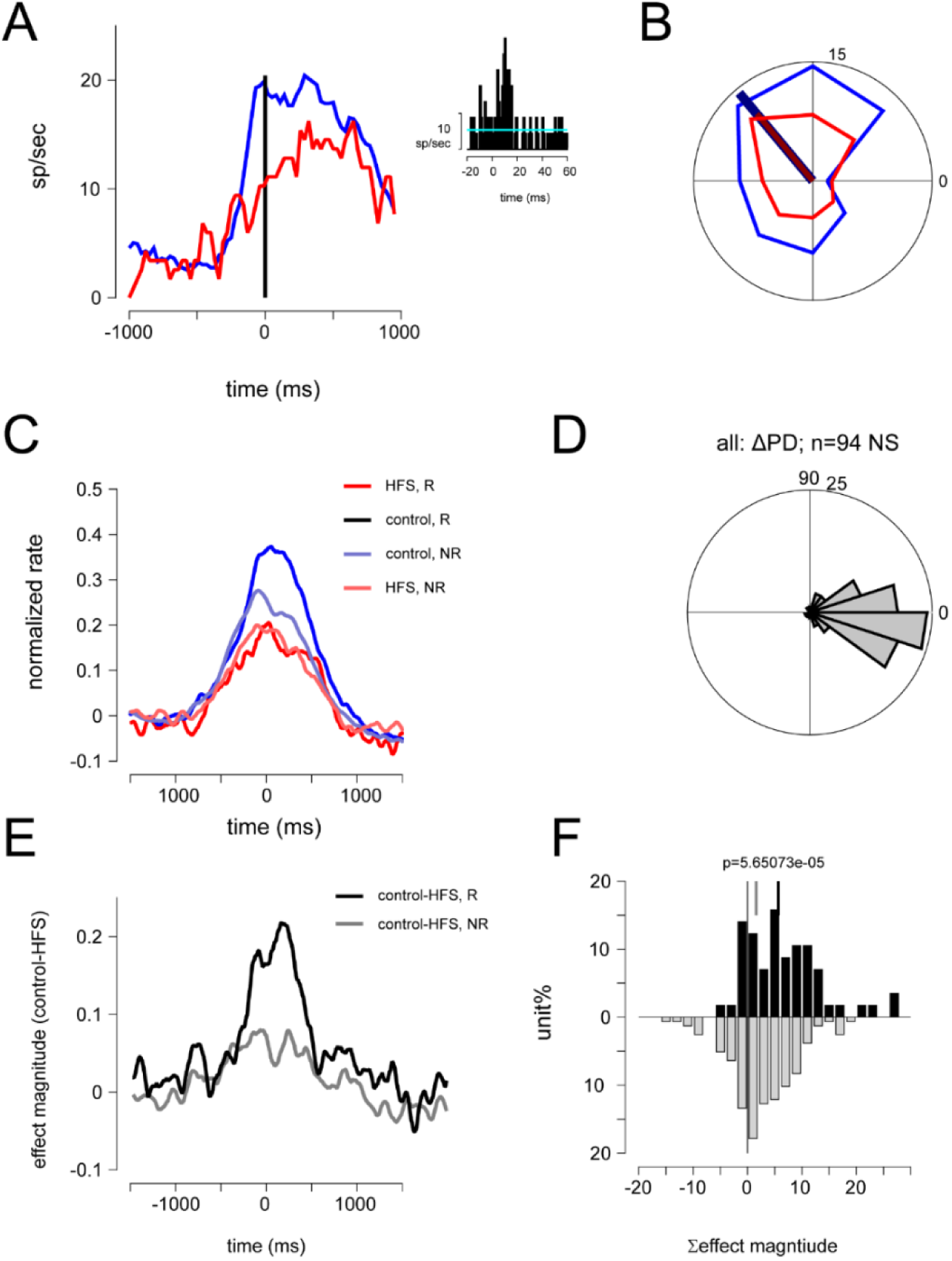
Task-related activity of cortical neurons was attenuated during HFS. (**A**) Example of task-related activity of an SCP-responsive neuron during control trials (blue) and HFS (red) trials around movement onset. Inset shows the response of the unit to single pulse SCP stimulation. (**B**) Tuning curve computed for the same neurons in (A) during control and HFS trials. The preferred direction of the cell is shown for both cases. (**C**) The mean normalized rate during control and HFS trials computed for responsive (R, n=57) or non-responsive (NR, n=158) neurons around movement onset. (**D**) Distribution of single-cell change in preferred direction between HFS and control trials (ΔPD) is shown for all cells that were tuned in both conditions. The mean value of the distribution was not significantly different from zero (one sample t-test for circular data with mean angle: p=0.59). The same results were obtained when considering only SCP responsive neurons (n=18, p=0.65 for responsive units). (**E**) Mean change in response profile computed by subtracting the single cell response obtained for HFS trials from the corresponding response for control trials and averaging the resulting function across all responsive (black) and non-responsive (gray)neurons. (**F**) For each neuron we quantified the change in response computed during HFS and control trials by integrating the normalized rate change over a time window spanning 50 to 350 ms around movement onset. The figure shows the distribution of these values computed for responsive (black) and non-responsive (gray) cortical neurons. The mean value of the two histograms are indicated by the vertical lines. The difference between these averages was significantly different (average accumulated change in responsiveness= 5.7±0.94 spikes, non-responsive=1.61±0.49 spikes, two-way t-test, p<0.001). **See also Fig. S3 and S4**.

If the phasic firing at movement onset is the result of the CTC drive, the SCP-responsive neurons should exhibit the greatest decrease in firing rate during HFS trials (compared to non-responsive cells). We tested this hypothesis and found that although both responsive (**Fig. 5C,E**) and non-responsive neurons reduced their transient firing rate around movement onset, the rate reduction observed for responsive units was significantly larger than for non-responsive cells (**Fig. 5F)**. This suggests that the effect of the HFS on behavior was predominantly mediated by the cortical neurons that are part of the CTC system. Importantly, it should be noted that the observed changes in task-related activity of the neurons recorded during HFS trials preceded the changes in muscle activity considerably (**Fig. S3**) such that the changes in neural activity could not simply be a reflection of the altered sensory feedback triggered by the impaired motor performance. We further confirmed that the changes in cortical activity and motor behavior were not specific to the behavioral paradigm used in this study. To do so, we applied HFS in a monkey that was trained to perform an isometric, single joint, delayed-response paradigm (Cohen et al., 2017; Nashef et al., 2018). HFS in this task affected motor behavior and cell activity in a manner similar to the findings reported here (**Fig. S4**), suggesting a general impact of HFS on movement-related activity, irrespective of the context in which movements are performed.

The impact of CTC system on movement onset raises the question of possible differences in CTC impact between the primary and premotor areas. Many studies have implicated the premotor areas in initiating movements (Kaufman et al., 2014; Mazurek and Schieber, 2017). Since HFS modified movement onset time considerably, we tested whether premotor neurons were uniquely affected by this manipulation. To do so, we divided the M1 and PM neurons that responded to SCP stimulation based on the peak time of their task-related activity into early (peak response before movement onset) and late (peak response after movement onset) cells (**Fig. 6A**). We found that although the majority of PM cells had an early peak response (as might be expected), fewer were responsive to SCP stimulation (**Fig. 6B**), compared to late PM cells (chi-square test, p <0.02). Moreover, the effect of HFS on early PM cells was considerably weaker compared to its effect on late PM cells (**Fig. 6C**). In contrast, for M1 neurons, both groups of cells (early and late) were comparably affected during the HFS trials both in terms of the fraction of SCP-responsive cells and the magnitude of HFS impact on their task-related activity (**Fig. 6B,D**). This suggests that the effect of HFS is not evenly distributed both in space (M1 vs. PM) nor in time; rather, the CTC system targets execution-related cells and its impact is strongest around movement onset.

**Figure 6.**
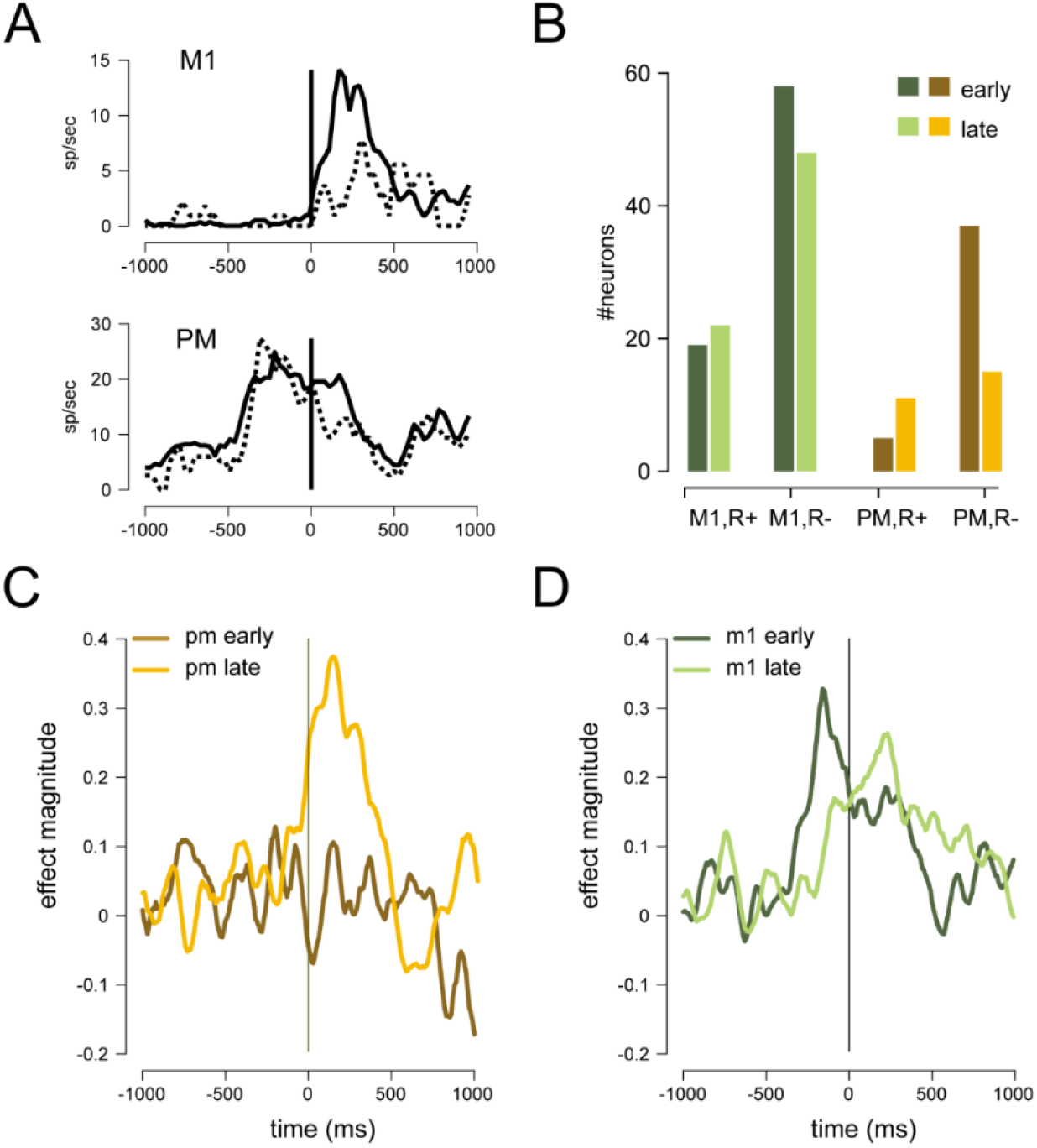
Differential effect of HFS on early and late motor cortical activity. (**A**) Example of a single cell response computed for an M1 neuron (upper panel) with a late peak response and a PM neuron (lower panel) with an early peak response. For both cells we show the average response during control (solid) and HFS (dashed) trials. Response peak time (early or late) was defined based on peak-time relative to movement onset during control trials. (**B**) Number of responsive (R+) and non-responsive (R-) neurons in M1 and PM areas segregated by response peak time (early – dark hues, late - light hues). For PM neurons, there were fewer early responsive neurons (n=5) than late responsive cells (n=11), whereas the reverse was true for non-responsive cells (early, n=37; late, n=15). These differences were significant (p < 0.02, X^2^ test). For M1 neurons there were no significant differences in early vs. late cells for responsive and non-responsive neurons (responsive: early=19, late-22; non-responsive: early= 58, late=48). (**C**) The single cell changes in response profile during HFS compared to control trials was computed, normalized and averaged separately for early and late PM neurons that responded to SCP stimulation. (**D**) same as (C) but for M1 neurons.

### Motor impairments during HFS trials are accompanied by local and global neuronal desynchronization

Applying HFS not only impeded movement onset but also impaired motor coordination. To identify the neural correlates of motor incoordination, we measured the neural interactions during the control and HFS trials. First, we estimated the noise correlation between simultaneously recorded neurons (**Fig. 7**). During the control trials, neighboring neurons (recorded by the same electrode) expressed a significant trial-to-trial rate co-variation (i.e., a positive noise correlation), especially when they shared similar tuning properties (i.e., had a positive signal correlation, **Fig. 7A**). This finding is similar to previous reports on motor cortical neurons (Lee et al., 1998). However, during HFS trials, the noise correlation between neighboring neurons decreased and no longer differed from zero (**Fig. 7B**); specifically, the firing of neighboring neurons became independent, despite the fact that the signal correlation remained the same (**Fig. 7C**). This result suggests that although the directional tuning of neurons was unaffected when flow of information in the CTC pathway was impaired, the local trial-to-trial synchrony was reduced in a way that was seemingly at odds with the increased behavioral variability characterizing HFS trials. Here again, the effect was stronger for responsive SCP than non-responsive neurons (**Fig. S5**).

**Figure 7.**
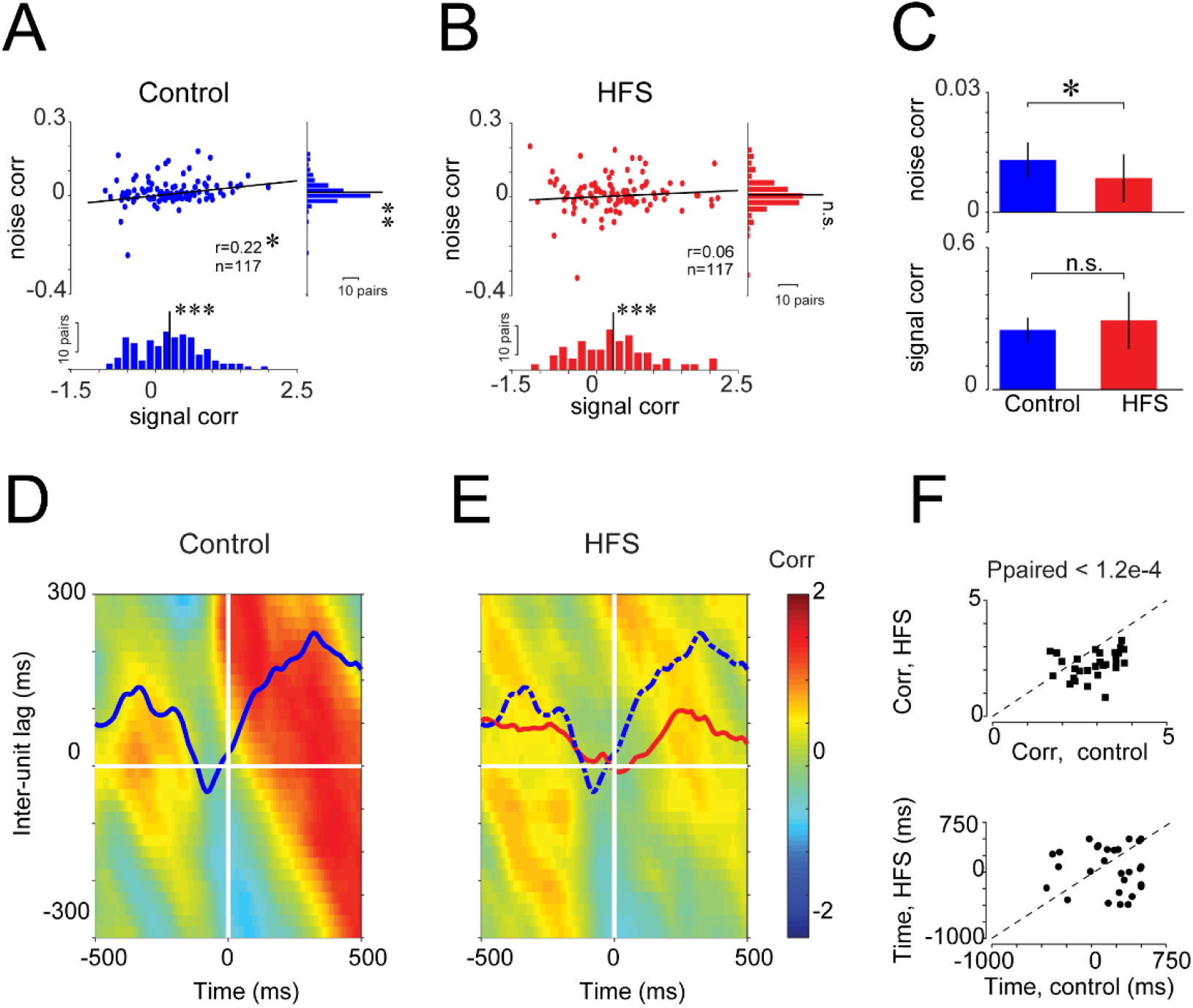
Reduced motor cortical synchrony during HFS. (**A**) Relation between noise and signal correlation for pairs of cortical neurons recorded from the same electrode (n=117 pairs) during control trials. The marginal distribution of each parameter and its deviation from zero are captured by the two histograms (P_noise_ < 0.005, P_signal_ < p<10^−5^ one-sample t-test). The correlation between noise and signal was tested as well (r=0.22, p < 0.05). (**B**) Same as (A) but for correlation values computed during HFS trials. For the marginal distribution, P_noise_ < 0.15 and P_signal_ < 10^−5^. The noise-signal correlation was r=0.06 (n.s.). Scale bars: 10 pairs. (**C**) The average noise (top) and signal (bottom) correlation during control and HFS trials (Kolmogorov-Smirnov test, p<0.05 for noise and n.s. for signal correlation). (**D**) Mean response correlation between pairs of neurons recorded simultaneously from shoulder-related and elbow-related cortical sites during control trials. The aligning event was the peak time of shoulder-related activity computed separately for each pair of neurons. Response correlation was measured for each time along the trial (x-axis) and inter-neuron time lag (y-axis). To average across all pairs, correlation matrices were transformed into Z-values. Bold line over the matrix represents the average correlation value computed for zero (inter-unit) lag. (**E**) Same as (D) but correlations computed during HFS trials. Here, the red curve depicts the mean correlation for the zero (inter-unit) lag during HFS trials, and the blue dashed curve is taken from panel (D) and is shown here for comparison. (**F**) Top: Pairwise comparison between peak correlation values for the time-resolved correlation during control (x-axis) and HFS (y-axis) trials. Peak values were computed for each pair of neurons using the time resolved correlation at zero inter-unit lag time (similar to the curves highlighted in panels (D,E)). Correlation values on control and HFS trials were slightly dependent (Spearman correlation coefficient = 0.38, p < 0.036) but significantly lower during HFS compared to control trials (p<0.00012, Wilcoxon signed-rank). Bottom: Same but for peak correlation time. There was no significant correlation between peak times in the control and HFS. **See also Fig. S5-7**.

Further, the loss of neural correlation also affected remotely located neurons. As proper performance of the task required shoulder-to-elbow coordination, we tested the specific relationships in responses between neurons recorded from shoulder and elbow-related sites during the control and HFS trials (**Fig. 7D-F**). Site identity was defined by the observed joint movement in response to intracortical microstimulation. We computed the pairwise response correlation between neurons recorded at elbow-related and shoulder-related sites. Since we searched for consistent temporal relations between activity patterns, for each pair of cells, we used the peak response time of the shoulder-related neuron (see Methods) as an aligning event when computing the peri-event time histogram (PETH) of the two cells. We then averaged the pairwise response correlation matrices across all available pairs. During the control trials (**Fig. 7D)**, we found that the response correlation varied along the trial, but was maximal at positive latencies (i.e., after the peak time of the shoulder-related cell). For each point along the trial, the maximal correlation was obtained at different shoulder-to-elbow time lags. For instance, at t=0 (peak shoulder activity) the maximal correlation was at positive time lags (i.e., when the elbow-related activity lagged about 200 ms behind the shoulder related activity). This suggests a consistent temporal organization of neural activity as concerns task-related joints (shoulder and elbow). By contrast, during HFS trials (**Fig. 7E)**, the temporal relations between the response profile of the shoulder and elbow units was lost both along the trial, and across different inter-unit lags. A quantitative examination of the changes in the pairwise response correlations revealed that during HFS, the value of the peak correlation decreased significantly (**Fig. 7F, top**, paired t-test, p<0.00012) and the peak time appeared more variable and uncorrelated with the peak time computed during control trials (**Fig. 7F, bottom**, Spearman correlation = −0.08, ns). For comparison, we computed the response correlation between pairs of neurons irrespective of their somatotopic affiliation (**Fig. S6A-C**). Here, there were no temporally organized activation patterns during the control or the HFS trials. The same result was obtained when computing the correlation between the shoulder-related neurons and neurons from task-irrelevant joints (i.e. wrist or fingers, **Fig. S6D-I)**, indicating that the observed temporal correlation of activity was highly dependent on the somatotopic identity of the tested neurons. Finally, we applied the same analysis, but using movement onset (instead of shoulder peak time) as an alignment event (**Fig. S7**). Here we obtained similar results, but the correlations values during the control trials were lower than those found when aligning on response peak time, indicating that the correlation matrices indeed measured the temporal locking of joint-related activity (which varied across trials in relation to movement onset). Taken together, these results suggest that in the absence of cerebellar input to the motor cortex, there is a spatiotemporal disorganization of motor cortical activity at the cell-to-cell level. This uncorrelated network activity is likely to underlie the uncoordinated movements that were observed during this time.

## DISCUSSION

The cerebellar-thalamo-cortical pathway (CTC) has long been considered a necessary component for executing well-timed and coordinated movements (Hore and Flament, 1988; Ivanusic et al., 2005; Proville et al., 2014). However, the mechanisms used by this system to control the properties of motor actions are not fully understood. Here we addressed this question by identifying the cortical components of the CTC system in behaving primates and by manipulating the flow of information through this pathway in a rapid but reversible manner. The behavioral consequences of this interference replicated symptoms of cerebellar ataxia, including longer response and movement times, curved movements and reduced inter-joint coordination. At the same time, neural activity was modified compared to the control trials, with a particularly pronounced suppression of task-related firing transients in the cortical neurons that were part of the CTC system and a loss of cell-to-cell synchrony in a manner consistent with the lack of motor coordination. The modification of cell activity was confined around movement execution, whereas the spatial tuning and early preparatory activity in premotor areas were unaffected. These results suggest that the CTC volley controls motor timing by synchronizing neurons both locally and globally. Local synchrony may contribute to the formation of firing transients which are effective in recruiting downstream elements and enhancing motor cortical throughput. The global synchrony which was found across task-related joints may support the appropriate temporal organization and the coupling of relevant effectors needed to execute specific motor tasks. Taken together, these actions of the CTC system facilitate rapid and coordinated motor execution, but are likely to have little role in the emergence of the motor plan.

Previous studies on the role of the CTC system in motor control have examined the functional implications of cerebellar deficits in human patients (Bastian et al., 1996; Bo et al., 2008; Deuschl et al., 2000; Spencer et al., 2003) or in primates where the dentate nucleus of the cerebellum was inactivated using a cooling probe (Beaubaton et al., 1978; Hore and Flament, 1988; Meyer-Lohmann et al., 1975). These studies provided invaluable information about the motor and neural consequences of cerebellar ataxia, but did not attempt to identify the set of neurons that constitute the CTC system or how the loss of the CTC drive affects single cell activity and leads to motor impairment. Here we utilized the anatomical organization of the CTC pathway and developed a new method to study the CTC system by using a chronically implanted SCP electrode. Targeting the SCP and not the cerebellar receiving motor thalamus provided a more specific access to the CTC pathway without recruiting neighboring thalamic nuclei. This type of confounding effect of intra-thalamic stimulation is a likely outcome, given the complex anatomy of the motor thalamus (Percheron et al., 1996). In fact, recent studies have suggested that the SCP is potentially a more efficient target for deep brain stimulation when treating essential tremor (Fenoy and Schiess, 2017).

The selected frequency of stimulation for the HFS was based on protocols commonly applied during stimulation of deep brain structures to treat neural disorders such as Parkinson’s disease (Benabid et al., 1991; Limousin et al., 1998). The effectiveness of the CTC block during HFS was clear from the fact that the cortical cells which were responsive to single pulse stimulation became non-responsive when the same stimulation pulses were applied at a high frequency. Since we stimulated a fiber tract, the block of information flow during HFS was probably due to the general inability of synaptic contacts to faithfully follow such repetitive activation (Wang and Kaczmarek, 1998; Zucker and Regehr, 2002). These findings are consistent with a recent report (Gornati et al., 2018) showing that the cerebello-thalamic transmission itself is blocked when activated at high frequencies. This suggests that high frequencies, SCP stimulation is an efficient method for blocking information transfer through the CTC system in a way which can be rapidly reversed when the stimulation is halted.

The behavioral consequences of the block of CTC information flow were similar to the symptoms of cerebellar ataxia. Cerebellar patients exhibit several stereotypical changes in motor behavior (Bastian et al., 1996; Diener and Dichgans, 1992; Holmes, 1939) including increased reaction time (Holmes, 1939; Schlerf et al., 2007; Tsujimoto et al., 1993), asymmetric movements (Diener and Dichgans, 1992; Holmes, 1939; Hore et al., 1991), decreased movement velocity (Bastian et al., 1996; Deuschl et al., 2000), end-point error and target overshooting (Bastian et al., 1996; Deuschl et al., 2000), decomposition of movement (Bastian et al., 1996; Becker et al., 1990) and tremor (Carrea and Mettler, 1955; Deuschl et al., 2000; Holmes, 1917, 1939). Motor coordination is specifically impaired in cerebellar patients (Bastian et al., 1996) as can be seen in the extensive deficits of these patients when performing multi-joint compared to single joint movements (Goodkin et al., 1993; Holmes, 1939; Thach et al., 1992). Most of these symptoms were replicated in the motor behavior of the monkeys during HFS trials; namely, response time and movement time increased, and movement velocity decreased. Since we used a shoulder-elbow reach paradigm we were able to test the impact of HFS on motor coordination. During HFS, the latency between the activation of the shoulder and elbow joints increased, suggesting a decomposition of movement. In addition, the synchrony between the shoulder and elbow joints was more variable and the movement became more curved. All these changes have been reported in cerebellar patients (Bastian et al., 1996; Becker et al., 1990; Deuschl et al., 2000).

The altered motor behavior observed during HFS trials was accompanied by substantial changes in neuronal firing patterns. One of the major changes in single cell activity during HFS was the loss of the phasic component of the response profile at movement onset. This change was consistent with the delayed onset and slower velocity of the action. Nonetheless, the change in firing considerably preceded muscle activity and movement. This suggests that the observed modification was in fact the source rather than the outcome of the poorly performed movements. Similar results have been found in studies that induced ataxic behavior using dentate cooling (Hore and Flament, 1988). Here we corroborate and extend these findings by specifically identifying motor cortical neurons that were part of the CTC system, as we first reported in a previous study (Nashef et al., 2018). The effect of rate change was particularly pronounced for these cells compared to non-responsive cells. This result directly implicates the CTC system in generating the firing transient at movement onset during normal motor behavior. This finding is also noteworthy as it indicates that the net impact of the TC input to the motor cortex exceeds the expected impact based on the low fraction of synaptic contacts made by the TC system on motor cortical cells (Bopp et al., 2017). Finally, despite the changes in response profile during HFS, the directional tuning of the cells remained the same. This strongly indicates that the spatial and temporal properties of motor cortical neurons are dictated by independent sources of information.

We also investigated the neural correlates of motor coordination (and its disintegration during HFS) at the level of the motor cortical network. The anatomy of the motor TC system seems to be specifically suited to coordinating motor actions (Horne and Butler, 1995) since a single TC fiber can generate multiple patches of terminals across several millimeters in the rostrocaudal axis (Shinoda et al., 1993). This contrasts with the spatially confined and locally dense distribution of TC terminals in the somatosensory system (Jones, 1983). These differences may reflect the different tasks and constraints the two systems face: whereas somatosensory information is transmitted in a way that preserves the somatotopic map, the motor TC system needs to coordinate several spatially distributed effectors during task performance. Our results highlight the possible neural correlates that correspond to the organization of the TC system and its contribution to motor coordination. Locally, we found that neighboring cells are correlated in a signal-dependent manner, consistent with previous reports (Lee et al., 1998). More globally, we found a temporal organization in the activation of joint-related sites. These two measures of neural coordination were lost during HFS, together with the deterioration in motor coordination.

In the motor cortex, the synchrony between neurons was reported to be linked with the similarity in directional tuning (Lee et al., 1998), shared muscle fields (Jackson et al., 2003) or specific motor states (Baker et al., 2001). All these findings appear to imply that neurons that share common features are likely to exhibit rate-based synchrony. The source for this rate covariation has not been explored directly, but it is assumed that either shared input and/or local recurrent interactions via intracortical axon collaterals are responsible for locally synchronized firing (Lee et al., 1998). The finding that during HFS the pairwise noise correlation between nearby neurons was by and large lost but the signal correlation was unaffected implies that at least part of this correlated noise among nearby units originates from CTC input which is independent of the directional tuning of the cells. The involvement of thalamic input in synchronizing the firing of cortical cells is the subject of considerable debate. In the somatosensory cortex, some have argued that common input may be sufficient to induce such synchrony (Bruno and Sakmann, 2006) whereas others have highlighted the role of intra-cortical processing for the observed synchrony (Cohen-Kashi Malina et al., 2016). Our results suggest that the interaction between these two sources of input (thalamic and cortical) is required for maintaining neural synchrony, such that in the absence of any of these inputs the cell-to-cell correlation is lost. The cortical synchrony supported by the TC system can account for the impact of the CTC system on cell firing, which extends beyond the anatomical connectivity alone.

In studying the relations between noise correlation and tuning similarity, a significantly positive noise correlation was only found between nearby neurons recorded by the same electrode (Lee et al., 1998). We speculate that documenting cells located at different patches that are formed along the trajectory of a single TC pathway would reveal remotely located cortical cells with positive noise correlations. We could not test this hypothesis directly, but we found that cortical cells that were recorded from the shoulder and elbow sites expressed a temporally organized response profile that was lost during HFS. Previous studies have reported large-scale propagating waves in the motor cortex that encode task parameters (Riehle et al., 2013; Rubino et al., 2006) and were related to motor initiation (Best et al., 2017). Studies have also shown that on average, the activation times of muscles and joint-related motor cortical cells occurs sequentially (from proximal to distal) with a large degree of overlap in onset times (Murphy et al., 1985). We extend this argument by suggesting that the temporal relations between remote sites that are related to different joints could be a way for the CTC system to control the timely activation required for multi-joint movements. In the absence of the CTC drive, these temporal relations are lost and the ability to properly coordinate different joints is impaired. Motor coordination was claimed to be the ability to exploit torques at one joint that were generated in moving another coupled joint (Bastian et al., 1996; Bastian et al., 2000). Hence, at the neural level, these torque interactions may be mediated by trial-to-trial synchronization across distinct populations of cells.

In summary, physiological and anatomical studies have shown that the CTC pathway is an extremely potent system with a broad motor cortical termination pattern. We showed that one of the functional implications of this organizational scheme is the synchronized recruitment of a motor cortical subnetwork composed of task related neurons that triggers and organizes the activation of its associated effectors in a timely manner. The synchrony produced by this pathway can potentially exert a strong impact on downstream elements. Previous studies have highlighted the importance of motor cortical synchrony for efficient recruitment of muscles (Baker et al., 2001; Jackson et al., 2003; Vaadia et al., 1995; Yanai et al., 2007). Our findings suggest that the CTC system may be an important cause and orchestrator of this synchrony.

## ACKNOWLEDGMENTS

This study was funded by the Israel Science Foundation (ISF-1787/13). Additional funding was received from the German Israeli Foundation (GIF, grant I-1224-396.13/2012), the Jerusalem Brain Center (AN) and through the generous support of the Baruch Foundation (YP).

## AUTHOR CONTRIBUTIONS

Y.P. conceived the study, A.N., O.C., Z.I. and R.H. performed the experiment, A.N. analyzed the data, Y.P. and A.N. wrote the manuscript.

## STAR★Methods

### RESOURCE TABLE

| Reagent or Resource | Source | Identifier |
| --- | --- | --- |
| Experimental model: Organisms/strains |  |  |
| *Macaca fascicularis* | The Hebrew University | N/A |
| Software and Algorithms |  |  |
| MATLAB 2016b | Mathworks | https://www.mathworks.com/ |
| Other |  |  |
| NSEX-100 | David Kopf instruments | http://kopfinstruments.com/ |

### CONTACT FOR RESOURCE SHARING

Further information and requests for resources and reagents should be directed to and will be fulfilled by the lead contact, Yifat Prut.

### EXPERIMENTAL MODEL AND SUBJECT DETAILS

This study was performed on two adult female monkeys *(Macaca fascicularis*, weight 4.5-8 kg). The care and surgical procedures of the subjects were in accordance with the Hebrew University Guidelines for the Use and Care of Laboratory Animals in Research, supervised by the Institutional Committee for Animal Care and Use.

### METHOD DETAILS

#### Behavioral task and electrophysiological recordings

Data were obtained from two *Macaca fascicularis* monkeys (females, 4.5-8 Kg). The monkeys’ care and surgical procedures were in accordance with the Hebrew University Guidelines for the Use and Care of Laboratory Animals in Research, supervised by the Institutional Committee for Animal Care and Use. The two monkeys were trained to sit in a primate chair, wear an exoskeleton (KINARM, BKIN technologies) and perform a planar, shoulder-elbow reaching task. In this task, the monkeys were instructed to locate a cursor within a central target. After 500 ms, a peripheral target (one of 8 evenly distributed targets) appeared and the monkey had to wait until the central target disappeared (“GO” signal) and reach the cued peripheral targets. If the monkey moved the cursor to the correct target within the predefined time limits it was rewarded with a drop of applesauce. To encourage the monkey to predict the timing of the “go” signal, we limited the total time it had to reach the peripheral target to 500 ms and inserted a 200 ms grace period before the GO signal. Onset of movement within this time frame did not abort the trial (**Fig. 1B**).

After training was completed, a recording chamber (21×21 mm or 27×27 mm) was attached to the monkeys’ skull above the hand-related area of the motor cortex in a surgical procedure under general anesthesia. After a recovery and re-training period, we recorded motor cortical activity extracellularly. During recording sessions, glass coated tungsten electrodes (impedance 300-800 kΩ at 1,000 Hz) were inserted through the chamber to different cortical sites, mostly in the primary motor cortex (M1). The signal obtained from each electrode was amplified (×10^4^), and bandpass-filtered online (300-6,000 Hz). The signal was then digitized (24 kHz) and saved to disk.

#### Insertion of stimulating electrode into the superior cerebellar peduncle (SCP)

To insert a chronic stimulating electrode into the ipsilateral SCP, we implanted a small chamber above the estimated insertion point and used a post-surgery MRI to plan the electrode trajectory. A bi-polar concentric electrode (NSEX100, David Kopf Instruments, impedance range of 30-60 kΩ) and the evoked intra-cortical responses to stimulation through the electrode were used to verify its location (Nashef et al., 2018; Ruach et al., 2015).

#### Mapping cortical areas

We used a set of up to 4 movable glass-coated tungsten electrodes (impedance 300-800 kΩ at 1 kHz) to record from motor and somatosensory areas of the cortex. For each recording site we mapped the motor response by observing the motor response evoked by intra-cortical microstimulation (train of stimuli applied at 333 Hz for 50 ms at intensities ≤ 60uA). A site for which an observable motor response was obtained at a threshold level ≤15μA was defined as the primary motor cortex (M1). A site that evoked a motor response at higher amplitudes and was located more than 3mm anterior to the central sulcus was defined as premotor (PM). Stimulation in somatosensory sites were located caudal to the CS and either required high stimulus intensities to produce a motor response or, more often, did not produce any motor response. Figure 1C presents the recording maps obtained for the two monkeys (2 hemispheres for monkey C and 1 for monkey M).

#### SCP stimulation protocol

To stimulate the SCP we used the following protocols.

##### Single pulse stimulation

Single stimulation pulses were applied via the SCP electrode while the monkey performed the task and neuronal activity was recorded. Each stimulation pulse was biphasic (200 μs each phase). A single set of stimuli consisted of about 200 stimuli that were delivered at 3 Hz and a fixed intensity (ranging from 50 to 300 μA).

##### High frequency stimulation (HFS)

We applied a long train of stimuli at high frequency through the SCP electrode during task performance and while recording cortical activity. HFS consisted of biphasic single pulse stimuli that were applied at 130 Hz and for a period of 120-180 seconds. Each train was delivered at a fixed intensity of 50-150 μA.

Each recording session (from a specific recording site) was usually tested in the following manner: (1) a set of control trials (about 80 trials); (2) 2-3 sets of single pulse stimulation applied at different intensities were applied; (3) pre-HFS control trials (~50 trials); (4) HFS trials lasting 120-180 seconds; (5) “washout” trials (with no stimulation). During this time the monkeys performed the task and neural activity was recorded.

#### EMG data

Muscle activity (EMG) was recorded from the two monkeys using transcutaneous electrodes inserted into selected arm and forearm muscles. The signals were filtered between 30 and 3,000 Hz and were digitized at 16 KHz per channel. EMG was recorded from the extensor-carpi-ulnaris, extensor-digitorum-carpi, extensor-carpi-radialis, flexor carpi ulnaris, palmaris longus, flexor carpi radialis, flexor digitorum superficialis, flexor digitorum profundus, extensor digitorum 2,3, abductor pollicis longus, brachioradialis, biceps, triceps, deltoid, and pectoralis major.

### QUANTIFICATION AND STATISTICAL ANALYSIS

All data were analyzed using MATLAB software (Mathworks).

#### Movement kinematics

During task performance we continuously measured the angular velocities of the shoulder and elbow joint and the endpoint position of the working arm. We used these measures to compute the following parameters:

1. Response time and movement time. Response time was calculated as the time between the appearance of the GO signal (the central target disappeared) and movement onset time. Since the monkey was allowed to move before the GO signal the response time was often negative, indicating a predictive timing control of the monkey. Movement time was defined as the time from movement onset until target acquisition (time in which the cursor entered the peripheral target).
2. Peak velocity. Maximal angular velocity (specified in degrees/second) of the working joints was calculated in a time window spanning −2 to 2 seconds around the GO signal.
3. Onset time of joint movements. The onset time of movement was defined separately for the shoulder and elbow joints in the following manner: for each trial, the single-joint radial velocity was first smoothed using a moving average spanning 10 bins. The baseline level and baseline noise (i.e., standard deviation around the mean – STD) of the joint velocity were calculated for a time window spanning −750 to −500 ms before the GO signal, when no movement occurred. We then focused on a time window starting at −250 before the GO signal, and searched for the time when the velocity signal deviated from the baseline level, and crossed a threshold of ± 20xSTD and remained above or below this threshold level for at least 200 ms. From this starting point, we went back to find the point where the velocity level crossed the baseline threshold ± 0.5xSTD. This point was defined as the onset time of joint movement.
4. Movement curvature. We calculated the average distance of the actual enacted trajectory from a direct line connecting the start and end points of the trajectory. This parameter was previously defined as the *curvature index* (Deuschl et al., 2000).

### Analysis of Electrophysiological Data

#### Stimulus-evoked responses of single neurons

The first step in the offline processing was to remove the stimulation artifacts from the neural signals by subtracting the average profile of the stimulation artifact from the raw signal (Ruach et al., 2015). Subsequently, an offline-sorting method (AlphaSort, Alpha-Omega, Nazareth, Israel) was applied on the cleaned signal to extract spike times of single units. We then calculated the SCP-evoked responses for each single unit by computing the peri-stimulus time histogram (PSTH) in a time window of −50 to +100 ms around stimulation time using a 1 ms bin size (Nashef et al., 2018). In short, background firing was computed from −50 to −10 ms before stimulus onset. The post-stimulation response was tested using two different time windows: strongly-locked early responses within a window of 1 to 8 ms using 1 ms bins were identified by t-testing the single trial spikes against the expected counts given the baseline rate level. A second more global test was then carried out in which we again tested the post-stimulus firing rate against background firing in a sliding window of 5 ms, shifted in 1 ms steps (1 bin). We used a t-test for the single sweep firing rates relative to the background firing and identified significant responses, defined as those that deviated from the background level with a probability of less than 0.01/n, where n was the number of bins (a Bonferroni correction to compensate for the fact that each bin was tested several times). We identified excitatory and inhibitory responses in a time frame of 2 to 45 ms by searching for at least 2 successive significant bins for excitatory responses or 7 significant bins for inhibitory responses.

#### Task-related response properties of recorded neurons

For each single unit, we computed the tuning function and its preferred direction (PD). The preferred direction was calculated individually for each isolated unit using a resampling method (Crammond and Kalaska, 1996; Shalit et al., 2012) (4,000 repetitions) during the - 500 to 500 ms around movement onset.

#### Estimating firing rate during HFS

The persistent high frequency stimulation produced many stimulus artifacts, some of which masked the action potentials emitted by the recorded neurons because the recorded signal was often saturated during part of the stimulus artifact. This random omission of spikes did not affect the response pattern of the neuron but yielded a lower estimate of its firing rate. To compensate for this loss of spikes when measuring the response pattern of each neuron we first counted the number of stimuli that were applied in each time bin of the peri-event time histogram (PETH) that was used to measure the neuronal response pattern. Normally, the firing rate for each bin is computed as the total number of spikes for that bin over the total time (Nspikes/T where T = bin width x number of trials). During HFS trials, we subtracted the time loss due to the post-stimulation dead time from T. This means that instead of T we used T − Nstim × 0.5 ms. The dead time was obtained by calculating the stimulus-triggered average of the single cell activity during HFS, which was found to be consistent across cells.

#### Measures of correlated firing between simultaneously recorded neurons

We used two measures to quantify the correlation in firing between pairs of cortical neurons that were recorded at the same time through the same or different electrodes.

#### Signal correlation

For each neuron, we computed the target-related signal (i.e., tuning curve) around movement onset (−500 ms before movement onset to +500 ms after). This tuning curve was simply the mean firing rate of the neuron while moving toward each target across trial repetitions. For each pair of simultaneously recorded neurons we computed the signal correlation by calculating the correlation coefficient between their target-related signals (Lee et al., 1998).

#### Noise correlation

For a single trial and a specific time window (−500 to +500 ms around movement onset) we computed the instantaneous noise by subtracting the average firing rate from the instantaneous firing rate, computed for trials with the same target during that time window. The noise correlation was then calculated as the correlation coefficient between the instantaneous noise of pairs of simultaneously recorded neurons. For each pair of neurons, we only considered targets for which we had at least 5 trials during the control and 2 during HFS. Furthermore, we only considered pairs of cells for which we had data for at least 10 trials. All the signal and noise correlation values were z-transformed to normalize their distribution in the following manner: *z* = 0.5 × [ln(1 + *r*) − ln(1 − *r*)].

The correlation values obtained for HFS trials relied on a smaller number of trials compared to the corresponding values obtained for the control trials. To compensate for this difference, we used a bootstrap approach and randomly selected a subset of control trials with a matching number of trials as obtained during HFS. The noise and signal correlation for the control trials was computed for this random subset. This procedure was repeated 1,000 times for each pair of neurons. The noise and signal correlation for control trials was then taken as the mean of the distributions.

#### Time-resolved response correlation of neural pairs

We examined the temporal relations between response profiles of simultaneously recorded neurons. This was done by first computing the PETH of the trigger neuron at its preferred direction (±1 target around the PD) around movement onset (±2,000 ms) and marking the time of peak activity. We then computed the PETH of a reference neuron aligned on the peak activity of the trigger neuron. The PETH was computed in a time window spanning −500 to +500 ms around the aligning event and in the preferred target ±1 target of the reference unit. To calculate the pairwise correlation matrix, we only considered neurons that had a roughly similar preferred direction (i.e., up, down, left or right). PETHs were computed using a 10 ms time bin.

We computed the time-resolved response correlation between pairs of neurons along trial time using a 200 ms time window shifted in 10 ms steps. For each point in time, the response correlation was measured between the two corresponding PETH vectors (each spanning 20 elements). We further computed correlation values after shifting the inter-unit time lags (−200…+200 ms in 10 ms steps). This means that each bin in the response correlation matrix corresponded to a specific time in the trial and a specific inter-unit time lag. Response correlation matrices were estimated for each pair of neurons and were subsequently z-transformed. Then, an averaged correlation matrix was computed by averaging across all matrices.

